# Detection of ctDNA from dried blood spots after DNA size selection

**DOI:** 10.1101/759365

**Authors:** Katrin Heider, Jonathan C. M. Wan, James Hall, Samantha Boyle, Irena Hudecova, Davina Gale, Wendy N. Cooper, Pippa G. Corrie, James D. Brenton, Christopher G. Smith, Nitzan Rosenfeld

**Affiliations:** Cancer Research UK Cambridge Institute, Li Ka Shing Centre, Robinson Way, Cambridge CB2 0RE, UK; Cancer Research UK Major Centre – Cambridge, Cancer Research UK Cambridge Institute, Li Ka Shing Centre, Robinson Way, Cambridge CB2 0RE, UK; Cambridge University Hospitals NHS Foundation Trust, Cambridge CB2 0QQ, UK

**Author notes:** K.H. and J.C.M.W. contributed equally to this work.

## Abstract

Recent advances in the research and clinical applications of circulating tumour DNA (ctDNA) is limited by practical considerations of sample collection. Whole genome sequencing (WGS) is increasingly used for analysis of ctDNA, identifying copy-number alterations, fragment size patterns, and other genomic features. We hypothesised that low-depth WGS data may be generated from minute amounts of cell-free DNA, and that fragment-size selection may be effective to remove contaminating genomic DNA (gDNA) from small volumes of blood. There are practical advantages to using dried blood spots as these are easier to collect, facilitate serial sampling, and support novel study designs in prospective human studies, animal models and expand the utilisation of archival samples by the removal of gDNA in small volumes. We therefore developed a protocol for the isolation and analysis of cell-free DNA from dried blood spots. Analysing a dried blood spot of 50μL frozen whole blood from a patient with melanoma, we identified ctDNA based on tumour-specific somatic copy-number alterations, and found a fragment size profile similar to that observed in plasma DNA processed by traditional methods. We extended this approach to detect tumour-derived cell-free DNA in a dried blood spot from a mouse xenograft model and were able to identify ctDNA from the originally grafted ascites. Together, our data suggests that ctDNA can be detected and monitored in dried blood spots. This will enable new approaches for sample collection from patients and *in vivo* models.

## Main text

### Introduction

Circulating tumour DNA (ctDNA) can be used to sensitively detect and quantify disease burden using a variety of sequencing based approaches^1^. For example, using shallow whole genome sequencing (sWGS), ctDNA can be detected down to mutant allele fractions of ∼3% through analysis of somatic copy-number alterations (SCNAs)^2,3^. Alternatively, leveraging differences in fragment size between tumour-derived and non-tumour cell-free DNA molecules (cfDNA) can enhance the detection of genomic alterations and the identification of plasma samples from patients with cancer compared to healthy individuals^4,5^. Although sWGS generates data on only a fraction of a single genome (0.3 genome equivalents correspond to ∼1 pg DNA), sequencing libraries for sWGS have traditionally been generated from larger amounts of cfDNA extracted from millilitre volumes of plasma from a venous blood samples^3^. Established protocols for collection of plasma for ctDNA analysis require prompt spinning of EDTA-containing tubes or delayed spinning of tubes containing cell preservatives/fixatives^6^. However, this processing restriction poses practical limitations on possible clinical study designs, especially in the context of serial sampling.

The use of limited blood volumes and dried blood spots for analysis of cfDNA may facilitate new trial designs, widen clinical applications, and enable point-of-care testing and self-collection of samples. Additionally, analysis of minute amounts of blood may facilitate longitudinal ctDNA monitoring from animal models with limited circulating blood volume. In prenatal diagnostics, polymerase chain reaction (PCR) has been used to carry out fetal *RHD* genotyping and HIV detection using maternal dried blood spots^7,8^. In applications to cancer, ctDNA from a limited plasma volume was previously analysed in a study of a mouse xenograft model, where quantitative PCR was used to measure the human long interspersed nuclear element-1 (hLINE-1) as a measure of tumour burden^9^. In another pilot study in breast cancer patients, whole genome amplification was performed on blood obtained from a finger prick. They found comparable allelic frequencies in somatic mutations between the finger prick sample and matched venous blood^10^. Sensitive detection of ctDNA from limited volumes or blood spots represents a technical challenge due to the limited total number of mutant molecules. Whole blood samples are considered inferior to carefully-collected plasma samples due to the presence of contaminating genomic DNA (gDNA) from lysed white cells in whole blood^1,11^, which dilutes tumour-derived ctDNA signal. In this study, we present methods for cfDNA extraction from dried blood spots and the subsequent analysis and detection of ctDNA.

### Results

We sought to assess the number of cfDNA genome copies that can be sequenced from a single blood drop or dried blood spot. Based on previous reports, the median concentration of cfDNA is approximately 1600 amplifiable copies per mL of blood for patients with advanced cancer^12,13^. This translates to approximately 80 copies of the genome as cfDNA in a blood drop/spot of 50μL. Assuming a yield in the range of ∼60%-80% in DNA extraction and efficiency of ∼15%-40% in generating a sequencing library, this is estimated to result in approximately 7x-25x representation of the genome in sequencing libraries prepared from cfDNA from a single blood drop. We therefore hypothesised that low-depth WGS of cfDNA can be attainable from a dried blood spot after removal of genomic DNA.

To test this hypothesis, we thawed frozen whole blood from a patient with Stage IV melanoma, and transferred 50μL to a Whatman FTA filter paper card. After drying the card for 15 minutes, we performed DNA extraction and library preparation from the dried blood spot. An overview of the workflow is shown in Fig. 1A. Quality control using capillary electrophoresis revealed contaminating gDNA, as indicated by an excess of large DNA fragments (Fig. 1B). cfDNA fragments typically display a characteristic fragmentation profile with a prominent peak at 166bp^4,14^. This peak was not observed, likely due to the low mass of cfDNA in the blood spot and the larger amounts of gDNA. To remove contaminating gDNA fragments (> 500bp in length), we applied a right-side size-selection using AMPure beads, at a bead-to-sample ratio of 1:1 (Methods). For this selection, beads are added to the sample and initially the DNA bound to the beads (carrying high molecular weight DNA) is removed while the supernatant containing low molecular DNA is retained. In a second step, additional AMPure beads are added (in a bead-to-sample ratio of 7:1) to capture all the remaining small-size fragments in the supernatant^15^. We generated a sequencing library from the size-selected DNA using the Thruplex Tag-Seq kit, and obtained a total of 232,107,928 sequencing reads (PE150; Illumina HiSeq4000; Fig. 1A).

**Fig. 1.**
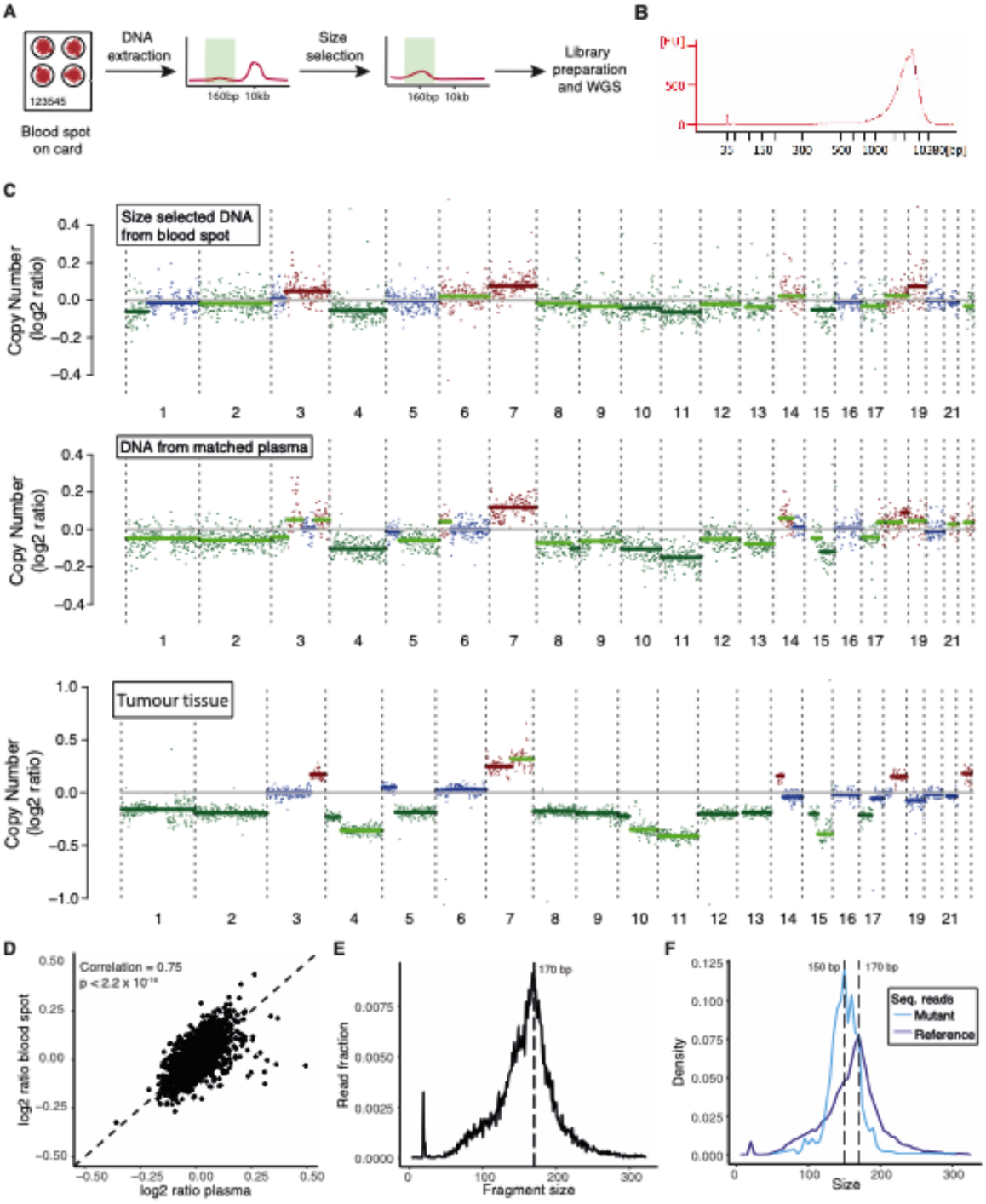
Detection of ctDNA in a dried blood spot from a cancer patient. (**A**) Overview of the analysis of dried blood spots: DNA extraction, followed by size selection, and low-depth WGS. (**B**) Bioanalyser trace of DNA extracted from a 50 µL dried blood spot from blood of a patient with advanced melanoma, showing a high level of genomic DNA contamination (> 1kbp) and no clear cfDNA peak (∼166bp). (**C**) Copy-number profiles from sWGS of a sequencing library generated from the same dried blood spot as in (B) after size selection, from a matched plasma sample from the same individual and timepoint and the matched tumour tissue. Blue=neutral, red=gain, green=loss. (**D**) Correlation of log2 ratios for each copy-number bin using iChorCNA^2^, comparing bins between matched blood spot and plasma data. The correlation in log2 ratios for all bins between the two samples was 0.75 (Pearson’s r, p < 2.2 × 10^−16^). (**E**) Size profile of the sequencing reads generated from the size selected blood spot DNA library (data shown in panel C). The overall size profile is comparable to that of cfDNA, i.e. with a peak at ∼170 bp. (**F**) Length of the sequencing reads (data from panel E) carrying known patient-specific mutations (light blue), and reads carrying reference alleles at the same loci (dark blue).

In our data, we achieved a unique sequencing depth of 6x from sWGS following collapsing with a minimum family size of 1^16^. Using a diversity estimator (SPECIES^17^), we inferred that, based on the distribution of the number of duplicate reads per molecule, up to 10x unique coverage are likely to be achieved from this blood spot library (Methods).

Sequencing data obtained from the blood spot was analysed for somatic copy-number alterations using ichorCNA^2^. The generated copy number plot is shown in Fig. 1C. The alterations observed were consistent with those identified in a matched plasma sample from the same patient, isolated by standard plasma DNA-based methods (Fig. 1C). The extent of SCNAs between the two samples was significantly correlated (Pearson’s r = 0.75, p < 2.2 × 10^−16^, Fig. 1D) and similar to that found in the initial tumour biopsy copy number profile (Fig. 1C).

Using sWGS, we show that the overall fragment size distribution of the human blood spot cfDNA was comparable to that of cfDNA derived from plasma^1,4,18^ (Fig. 1E). We then independently analysed the size distribution of mutant and wild-type reads, leveraging mutation calls from exome sequencing of matched tumour tissue in order to accurately distinguish true mutations from sequencing noise. This confirmed that the tumour-derived fragments were shorter in size compared to wild-type fragments, with modal sizes of 150 bp and 170 bp, respectively (Fig. 1F). These data recapitulate size profiles derived from plasma samples of cancer patients^1,4,18^.

We next considered whether blood spot analysis may have applications in the longitudinal analysis of disease burden in live murine patient derived xenograft (PDX) models. At present, analysis of cfDNA is challenging in small rodents as the volumes of blood required for most traditional ctDNA analysis can only be obtained through terminal bleeding. To assess the feasibility of dried blood spot analysis in animal models, we sampled 50μL of whole blood onto a dried blood spot card from an orthotopically implanted ovarian tumour PDX model. DNA was extracted and sequenced (Methods). Following alignment of sequencing reads, both human genome (tumour-derived) and mouse genome (wild-type) reads were observed, again showing characteristic fragmentation patterns of mutant and wild-type cfDNA^4^ (Fig. 2A). Copy-number alterations were observed when analysing the human sequencing reads and mirrored the profile observed in both the original patient ascites sample and the matched PDX tumour in the mouse (Fig. 2B). This confirms that blood spots can indeed be used to monitor disease progression and burden in animal models.

**Fig. 2.**
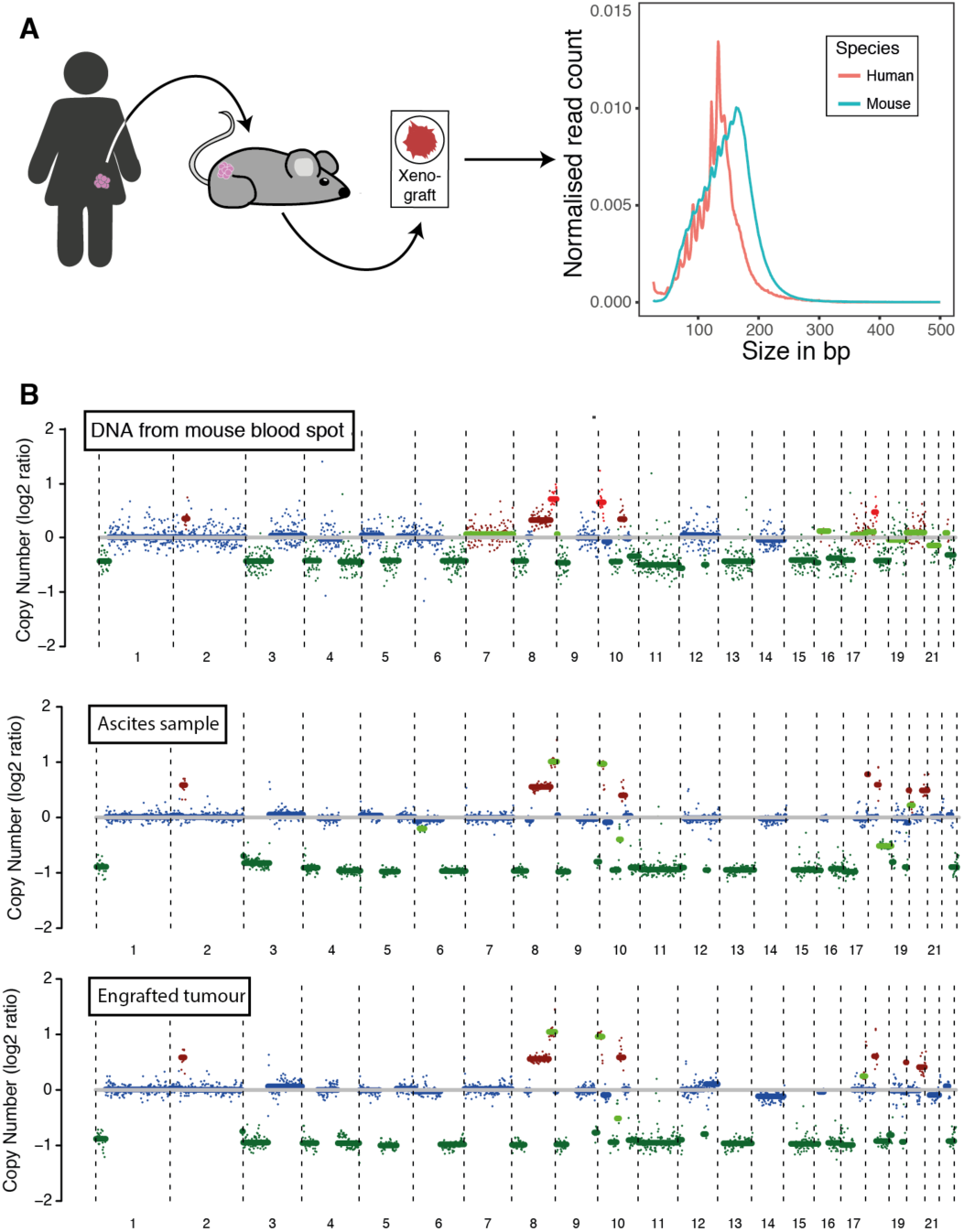
ctDNA detection from a dried blood spot in a xenograft model. (**A**) sWGS analysis of whole blood taken from a mouse xenograft model of ovarian cancer (illustrated in the left panel). The fragment lengths of reads aligning to the human genome (red, representing tumour ctDNA) were shorter than those aligning to the mouse genome (blue, representing non-tumour cfDNA). (**B**) Copy-number profiles were successfully generated from a dried blood spot from the mouse ovarian xenograft model (Methods). The copy-number profiles of the original human ascites sample and the engrafted tumour are also shown. Segments coloured in blue, red and green indicate regions of copy-number neutrality, gain and loss, respectively.

### Discussion

In this study, we demonstrate a new method to detect ctDNA in blood drops/spots using sWGS from both human and PDX samples. Our analysis mirrors observations previously made in cfDNA plasma analysis. This approach relies on the use of size-selection to remove genomic DNA, combined with ctDNA measurement approaches such as sWGS which leverage signal from across the entire genome. Only highly multiplexed approaches leveraging signal from multiple loci are suitable for blood spot analysis. The analysis of any individual locus would have limited sensitivity due to the small number of genome copies of cfDNA that may be obtained from a single blood spot (in the order of 5-50 copies).

We analysed a dried blood spot from a patient with melanoma and observed a good correlation in the copy-number profiles obtained from the blood spot and a time-matched plasma and tumour sample. We see similar cfDNA and ctDNA size profiles as observed from standard plasma DNA-based methods. Further work on larger cohorts with fresh finger prick blood is warranted before progressing towards broader use of blood spots for ctDNA monitoring. This work should compare the ctDNA allele fractions between blood spot and matched plasma, as the extent of gDNA contamination of blood spot DNA might vary. Additionally, the sensitivity limit for ctDNA analysis in blood spots should be determined with both sWGS and targeted sequencing approaches. If single nucleotide variants were to be targeted, a larger number of patient-specific mutations must be identified and interrogated in order to adequately mitigate the effect of sampling error from the limited copies of cfDNA in small volumes of blood. In future, the potential application of personalised sequencing panels^19^ to sequencing data could facilitate highly sensitive monitoring of disease from even small volumes.

In addition, we demonstrate the value of this approach in animal models, allowing the detection of SCNAs and the characteristic ctDNA fragmentation pattern from dried blood spots of PDX models. In the monitoring of ctDNA in small animal models, overcoming low circulating blood volumes is a major challenge. Although tail vein blood sampling in rodents has already been used for longitudinal cancer monitoring from small blood samples, analysis was limited to high copy-number markers such as hLINE repeat sequences^9^. Here, we highlight the possibility of next generation sequencing of blood spot cfDNA, enabling both, shallow and up to 10× WGS.

From a practical standpoint, the application of dried blood spots could enable high-frequency ctDNA monitoring of patients and animal models. Sampling and pre-analytical processing can be further simplified, potentially supporting new study designs incorporating wider populations and more frequent collection of smaller sample volumes. We hope that detection of ctDNA from limited blood volumes will enable novel approaches for cancer monitoring, such as self-collection of samples at home followed by shipping and centralised analysis.

## Methods

### Cell-free DNA extraction from dried blood spots

The human sample was collected from a patient enrolled on the MelResist study (REC 11/NE/0312). Written consent to enter the study was taken by a research/specialist nurse or clinician who was fully trained regarding the research. MelResist is a translational study of response and resistance mechanisms to systemic therapies of melanoma, including BRAF targeted therapy and immunotherapy, in patients with stage IV melanoma. Thawed whole blood (50 μL) was transferred to Whatman FTA™ Classic Cards (Merck) and allowed to air dry for 15 minutes before DNA extraction. A single fresh blood spot was obtained from an ovarian cancer xenograft mouse model immediately after culling, and similarly applied to Whatman FTA™ Classic Cards, and allowed to air dry. Blood spot card samples were stored at room temperature inside a re-sealable plastic bag. DNA was extracted from the card using the QIAamp DNA Investigator kit (Qiagen), using the manufacturer’s recommended extraction protocol for FTA and Guthrie cards, which are conventionally used for assessment of inherited genetic conditions in neonates from gDNA. The protocol was followed with the following modifications. 1) three 3mm punches were made from the blood spot, and carrier RNA was added to Buffer AL as per the manufacturer’s recommendation. 2) blood spot DNA (which we hypothesised contained both cfDNA and gDNA) was eluted in 25μL water, which was reapplied to the membrane and re-eluted.

### Size-selection and library preparation of blood spot cell-free DNA

Blood spot DNA eluates contain a low concentration of cfDNA, among a large background of gDNA (Fig. 1B). cfDNA library preparation cannot be effectively performed from such a sample since the abundance of long fragments reduces the likelihood of any cfDNA fragments successfully being ligated with adaptor molecules for subsequent amplification. Based on our characterisation of gDNA length of >1-10kb (Fig. 1B), and previous work demonstrating that cfDNA *in vitro* ranges from ∼70-300bp in length with a peak at 166bp^20^, we opted to perform size-selection in order to remove contaminating long gDNA fragments. Thus, a right-side size-selection was performed on DNA eluates using AMPure XP beads (Beckman Coulter) in order to remove long gDNA fragments. For this purpose, we adapted a published protocol for a right-side size selection that is conventionally used for DNA library size-selection prior to next generation sequencing^15^. Following optimisation of bead:sample ratios for cfDNA fragment sizes, we used a bead:sample ratio of 1:1 to remove contaminating gDNA. The supernatant was retained as part of the right-side size-selection protocol. A second size-selection step used a 7:1 bead:sample ratio to capture all remaining fragments, and the size-selected DNA was eluted in 20μL water. Blood spot eluates were concentrated to 10μL volume using a vacuum concentrator (SpeedVac), since this volume is the maximum recommended for downstream library preparation using the Thruplex Tag-Seq kit (Takara). 16 cycles of library amplification were carried out. Libraries underwent QC using Bioanalyser 2100 (Agilent) and qPCR with the Illumina/ROX low Library Quantification kit (Roche) on a QuantStudio 6 (Life Technologies). Libraries were submitted for whole-genome sequencing on a HiSeq4000 (Illumina) with paired end 150bp/cycles.

### Plasma library preparation

Plasma cfDNA libraries were prepared for the matched timepoint where the blood spot was collected as well as a cohort of 49 healthy controls. The DNA was extracted using the QIAsymphony (Qiagen) with the QIAamp protocol and quantified by digital PCR on a Biomark HD (Fluidigm) using a 65bp TaqMan assay for the housekeeping gene RPP30 (Sigma Aldrich)^21^ and 55 cycles of amplification. Using the estimated number of RPP30 DNA copies per μL eluate, the cfDNA concentration in the original sample was estimated. Up to 9.9ng were used for the library preparation. The ThruPLEX Tag-Seq kit (Takara) was used according to the manufacturer’s instructions and 7 cycles of amplification were carried out. After barcoding and sample amplification, the library underwent bead clean-up and underwent QC as described above. The sample was submitted for sequencing on a HiSeq4000 with paired end 150bp/cycles.

### Tumour library preparation

For the human blood spot, a time matched tumour sample was available. Tumour DNA was extracted as described by Varela et al.^22^ and sheared to ∼200bp fragment length using the COVARIS LE220 Focused-ultrasonicator according to manufacturer’s instructions. 50ng of material were prepared for sWGS using the ThruPLEX Plasma-Seq kit (Takara) according to the manufacturer’s instructions and 7 cycles of amplification were carried out. After barcoding and sample amplification, the library underwent bead clean-up and underwent QC as described above. The sample was submitted for sequencing on a HiSeq4000 with 150bp/cycles.

For the xenograft sample, material from the engrafted tumour as well as the human ascites sample used for grafting were available for analysis. The sample was extracted using the Qiagen allprep kit (Qiagen) and the DNA was sheared to 200bp fragment as described above. 50ng of DNA were prepared with the Thruplex DNA-Seq kit (Takara) according to the manufacturer’s instructions and followed by a bead clean-up (1:1 ratio, as described above). The sample was quantified using TapeStation (Agilent) and submitted for sequencing on a HiSeq4000 with single end 50bp/cycles.

### Sequencing data analysis

All samples were sequenced on a HiSeq4000. FASTQ files were aligned to the UCSC hg19 genome using BWA-mem v0.7.13 with a seed length of 19, then deduplicated with MarkDuplicates. For sWGS detection of ctDNA, iChorCNA was run as described^2^, utilising a panel of normal derived from a set of plasma cfDNA samples from 49 healthy individuals (SeraLabs).

For xenograft sequencing analyses, BAM files underwent alignment to the mouse and human genomes in parallel using Xenomapper^23^. Fragment lengths were determined for both files using Picard CollectInsertSizeMetrics^24^. Additionally, iChorCNA^2^ was run on the subset of reads aligning to the human genome to confirm the presence of CNA.

### Library diversity estimation

In order to estimate the total number of cfDNA genome copies present in a blood spot library, we used CONNOR^16^ to perform deduplication of the blood spot sequencing library based on endogenous barcodes^25^ with minimum family sizes ranging between 1 and 5 (data not shown). For each family size setting, the mean deduplicated coverage was calculated using Samtools mPileup. Deduplicated coverage values for each setting were used as input for diversity estimation using a statistical method, SPECIES^17^, best known for estimating the diversity of ecological populations based on the frequency of members observed through a random sample. A minimum family size of 1 was used for the data analysis.

## Author contribution

K.H., J.C.M.W. and N.R. wrote the manuscript. K.H. and J.C.M.W. carried out the experiments and data analysis. J.H. and S.B. generated and prepared the animal model. I.H. and W.N.C. helped in designing the study. P.G.C., N.R. and D.G. led and coordinated work on the MelResist study from which the human blood spot was used. C.G.S., J.D.B. and N.R. supervised the project.

## Acknowledgments

The authors would like to thank Catherine Thorbinson, Emily Barker and Alex Azevedo from the MelResist study group and the Cambridge Cancer Trials Centre, Addenbrookes Hospital, Cambridge. We also thank Carolin Sauer for acquisition of mouse samples and preparation of sequencing data.

